# Optical genome mapping enables accurate repeat expansion testing

**DOI:** 10.1101/2024.04.19.590273

**Authors:** Bart van der Sanden, Kornelia Neveling, Syukri Shukor, Michael D. Gallagher, Joyce Lee, Stephanie L. Burke, Maartje Pennings, Ronald van Beek, Michiel Oorsprong, Ellen Kater-Baats, Eveline Kamping, Alide Tieleman, Nicol Voermans, Ingrid E. Scheffer, Jozef Gecz, Mark A. Corbett, Lisenka E.L.M. Vissers, Andy Wing Chun Pang, Alex Hastie, Erik-Jan Kamsteeg, Alexander Hoischen

## Abstract

Short tandem repeats (STRs) are amongst the most abundant class of variations in human genomes and are meiotically and mitotically unstable which leads to expansions and contractions. STR expansions are frequently associated with genetic disorders, with the size of expansions often correlating with the severity and age of onset. Therefore, being able to accurately detect the total repeat expansion length and to identify potential somatic repeat instability is important. Current standard of care (SOC) diagnostic assays include laborious repeat-primed PCR-based tests as well as Southern blotting, which are unable to precisely determine long repeat expansions and/or require a separate set-up for each locus. Sequencing-based assays have proven their potential for the genome-wide detection of repeat expansions but have not yet replaced these diagnostic assays due to their inaccuracy to detect long repeat expansions (short-read sequencing) and their costs (long-read sequencing).

Here, we tested whether optical genome mapping (OGM) can efficiently and accurately identify the STR length and assess the stability of known repeat expansions. We performed OGM for 85 samples with known clinically relevant repeat expansions in *DMPK*, *CNBP* and *RFC1*, causing myotonic dystrophy type 1 and 2 and cerebellar ataxia, neuropathy and vestibular areflexia syndrome (CANVAS), respectively. After performing OGM, we applied three different repeat expansion detection workflows, *i.e.* manual *de novo* assembly, local guided assembly (local-GA) and molecule distance script of which the latter two were developed as part of this study. The first two workflows estimated the repeat size for each of the two alleles, while the third workflow was used to detect potential somatic instability. The estimated repeat sizes were compared to the repeat sizes reported after the SOC and concordance between the results was determined.

All except one known repeat expansions above the pathogenic repeat size threshold were detected by OGM, and allelic differences were distinguishable, either between wildtype and expanded alleles, or two expanded alleles for recessive cases. An apparent strength of OGM over current SOC methods was the more accurate length measurement, especially for very long repeat expansion alleles, with no upper size limit. In addition, OGM enabled the detection of somatic repeat instability, which was detected in 9/30 *DMPK*, 23/25 *CNBP* and 4/30 *RFC1* samples, leveraging the analysis of intact, native DNA molecules. In conclusion, for tandem repeat expansions larger than ∼300 bp, OGM provides an efficient method to identify exact repeat lengths and somatic repeat instability with high confidence across multiple loci simultaneously, enabling the potential to provide a significantly improved and generic genome-wide assay for repeat expansion disorders.

## INTRODUCTION

Short tandem repeats (STRs) are common repeats of a particular *k*-mer of 1-6 base pairs in length (Tankard et al. 2018). More than a million cataloged STR loci make up ∼3% of the human genome and are scattered throughout (Lander et al. 2001; Gymrek 2017). Expansions or contractions of at least 60 of these STRs have been associated with human genetic disorders, concerning predominantly neurogenetic diseases (Depienne and Mandel 2021; Tanudisastro et al. 2024). These disorders include, but are not limited to, myotonic dystrophies, Huntington disease, fragile X syndrome, and different forms of spinocerebellar ataxias (van der Sanden et al. 2021). STR disorders present with overlapping clinical phenotypes, strong heterogeneity of symptoms, and variation in age of onset, which makes identification of the molecular diagnosis challenging (Tankard et al. 2018).

All individuals have a certain repeat length at each disease-associated STR locus, however only once the size of a disease-associated repeat exceeds a certain repeat size threshold, the individual may develop a disorder. For several STR disorders a strong correlation between the size of the expansion and the severity as well as the age of onset of the disorder have been associated (Paulson 2018; Depienne and Mandel 2021). An important characteristic of dominant STR expansion disorders is anticipation, a phenomenon where new generations are affected at an earlier age of onset and with more severe symptoms than the preceding generations. In addition to anticipation, repeat expansions can present with somatic instability, a dynamic process in which the repeat size can increase over time, which may be tissue dependent (Monckton et al. 1995; Wong et al. 1995; Gomes-Pereira et al. 2004). For some repeat expansion disorders the disease severity increases when the repeat expansion is somatically unstable (Gomes-Pereira et al. 2004; Swami et al. 2009; Goold et al. 2021; Ruiz de Sabando et al. 2024). Finally, repeat expansions can contain interruptions – for example a CCG interruption in a CTG repeat expansion in DMPK – and these may cause a repeat expansion to be more stable than uninterrupted repeat expansions, thereby reducing somatic instability and leading to milder symptoms (Cumming et al. 2018; Nolin et al. 2019; Depienne and Mandel 2021). However, repeat expansions are largely heterogeneous and not all repeat expansion loci are equally affected by repeat interruptions or somatic instability.

The current standard of care (SOC) for patients with a suspected repeat expansion disorder can be time consuming and costly. The clinician must request the appropriate repeat expansion test based on the patient’s disorder. The standard of care then consists of targeted PCR and repeat-primed PCR (RP-PCR) and/or Southern-blot assays. These assays must be refined for each different repeat expansion locus, which means that the same sample may have to undergo multiple rounds of diagnostic testing. This can be due to phenotypic overlap between expansions of different STRs, heterogeneity of symptoms, and variation in penetrance and age of onset (Tankard et al. 2018). Over the last decade, exome sequencing (ES) has become increasingly important for diagnosing patients (Srivastava et al. 2019) and in addition to the targeted repeat expansion assays, it is now also possible to detect specific STR expansions using ES and genome sequencing (GS) (Gymrek et al. 2012; Dolzhenko et al. 2017; Tang et al. 2017; Willems et al. 2017; Dashnow et al. 2018; Tankard et al. 2018; Dolzhenko et al. 2019; Mousavi et al. 2019; van der Sanden et al. 2021). However, dedicated next generation sequencing STR detection tools are limited by the 100-150 bp read length and/or total fragment length of short-read sequencing (Halman and Oshlack 2020; Tanudisastro et al. 2024). Altogether, every genetic diagnostic test that is currently performed for patients with a suspected repeat expansion disorder has its own limitations and no generic one-test-fits-all approach is currently available.

The introduction of long-read technologies has allowed the detection of large repeat expansions and determining the exact repeat size because long reads can entirely span (very long) repeat loci, which improves mapping quality and reduces mapping bias (Mantere et al. 2019; Tanudisastro et al. 2024). Recently, long-read sequencing technologies, such as HiFi (PacBio) and nanopore (ONT) sequencing, have proven the benefit of long reads for STR detection (Giesselmann et al. 2019; Mitsuhashi et al. 2019; Sone et al. 2019; Chiu et al. 2021; Dolzhenko et al. 2024). However, the current high cost of long-read genome sequencing limits the widespread use of the technology for STR expansion detection (Tang et al. 2017). Therefore, targeted long-read sequencing approaches are emerging (Loose et al. 2016; Höijer et al. 2018; Miyatake et al. 2022; Stevanovski et al. 2022). Optical genome mapping (OGM) is another long-read technology, which generates images of ultra-long high molecular weight (UHMW) DNA molecules with average N50 > 250kb (Neveling et al. 2021). OGM has proven to provide a cost-effective and easy-to-use alternative for structural variant (SV) detection and is also capable of detecting STRs (Mantere et al. 2021; Neveling et al. 2021; Facchini et al. 2023; Guruju et al. 2023). In addition, OGM is independent of sequence context and in combination with the ultra-long molecules and genome-wide coverage it enables the analysis of even the most complicated regions of the genome in contrast to DNA sequencing approaches (Neveling et al. 2021). Therefore, OGM has a great potential for determining the exact repeat sizes of even the longest repeats.

In this study, we tested whether OGM can efficiently and accurately identify the repeat length across multiple STR loci simultaneously, thereby detecting large STR expansions and determining their absolute repeat sizes as well as potential somatic instability. To this end, we performed OGM for 85 samples with known clinically relevant repeat expansions in *DMPK*, *CNBP* and *RFC1* causing myotonic dystrophy type 1 and 2, and cerebellar ataxia, neuropathy and vestibular areflexia syndrome (CANVAS), respectively. Next, the OGM data was sequentially used in three different workflows. Firstly, the regularly available standard analysis workflow referred to as ‘manual *de novo* assembly’, secondly a local guided assembly and thirdly a molecule distance script. The latter two were developed and applied as part of this study. The first two workflows were used to determine the repeat size of both alleles, while the third workflow was mainly used to identify potential somatic instability. This approach allowed for a direct comparison of the repeat sizes estimated by OGM and the repeat sizes reported after the SOC, providing an evaluation of OGM as a repeat expansion detection technology.

## METHODS

### Patient selection

The department of Human Genetics of the Radboudumc is a referral center for patients with suspected repeat expansion disorders. In total, 85 patients with a known (bi-allelic) repeat expansion in *CNBP* (n=25), *DMPK* (n=30) and *RFC1* (n=30) were selected from our patient cohort and anonymized for further use in this study. Further repeat expansion details can be found in **Table 1**. This study was approved by the Medical Review Ethics Committee Arnhem-Nijmegen under 2011-188 and 2020-7142.

**Table 1:** Repeat expansion details.

| Gene | Disease | Inheritance | Location of repeat in gene | Repeat unit | Normal repeat size | Premutation repeat size | Pathogenic repeat size |
| --- | --- | --- | --- | --- | --- | --- | --- |
| *CNBP* | DM2 | AD | Intron | CCTG | <27 | 27-74 | >74 |
| *DMPK* | DM1 | AD | 3'UTR | CTG | 5-35 | 36-49 | >49 |
| *RFC1* | CANVAS | AR | Intron | AAGGG (pathogenic)<br>AAAAG (benign)<br>AAAGG (benign) | 11 (AAAAG) | n/a | >400<sup>#</sup> |

<sup>#</sup> SOC for *RFC1* repeat expansions is not suited to detect full repeat sizes. It uses a combination of locus-spanning PCR, resulting in allelic dropouts for repeats >120 units, and repeat-primed PCR to detect the repeats up to 20 units. For the sake of this technical study, repeat sizes >20 units were already considered expansions irrespective of their pathogenicity. Table adjusted from van der Sanden *et al.* 2021 (van der Sanden *et al.* 2021).

### Standard of care tests

PCR and fragment-length analysis, repeat-primed PCR and Southern blotting for *CNBP* and *DMPK* repeat expansions were previously performed as part of routine diagnostic repeat expansion testing according to previously described standard protocols (Kamsteeg et al. 2012). Locus-spanning PCR and repeat-primed PCR for *RFC1* repeat expansions were also performed as part of routine diagnostic repeat expansion testing according to previously described standard protocol (Ghorbani et al. 2022).

### DNA isolation, labeling and optical genome mapping

DNA isolation, labeling and optical genome mapping were performed as described previously (Mantere et al. 2021; Neveling et al. 2021). For each individual, UHMW DNA was isolated from 650 µL of whole peripheral blood (EDTA) or 1–1.5 million cultured cells using the SP Blood and Cell Culture DNA Isolation Kit according to the manufacturer’s instructions (Bionano, San Diego, CA, USA). Briefly, cells were treated with a lysis-and-binding buffer (LBB) to release UHMW DNA, which was then bound to a nanobind disk, washed, and eluted in the provided elution buffer. UHMW DNA molecules were labeled with the DLS (Direct Label and Stain) DNA Labeling Kit (Bionano, San Diego, CA, USA). Direct Label Enzyme (DLE-1) and DL-green fluorophores were used to label 750 ng of UHMW DNA. After a wash-out of the DL-green fluorophore excess, the DNA backbone was counterstained overnight before quantitation. Labeled UHMW DNA was loaded on a Saphyr® chip G2.3 for linearization and imaging on the Saphyr® instrument (Bionano, San Diego, CA, USA).

### OGM repeat expansion workflows

The entire data analysis was performed as previously described (van der Sanden et al. 2024). In the following section we only summarized the most important steps in the data analysis process.

The BNX molecule files generated by the Bionano Saphyr® machine were sequentially used in three different workflows (**Fig. 1**).

1. Manual *de novo* assembly
2. Local guided assembly (local-GA)
3. Molecule distance script

**Fig. 1:**
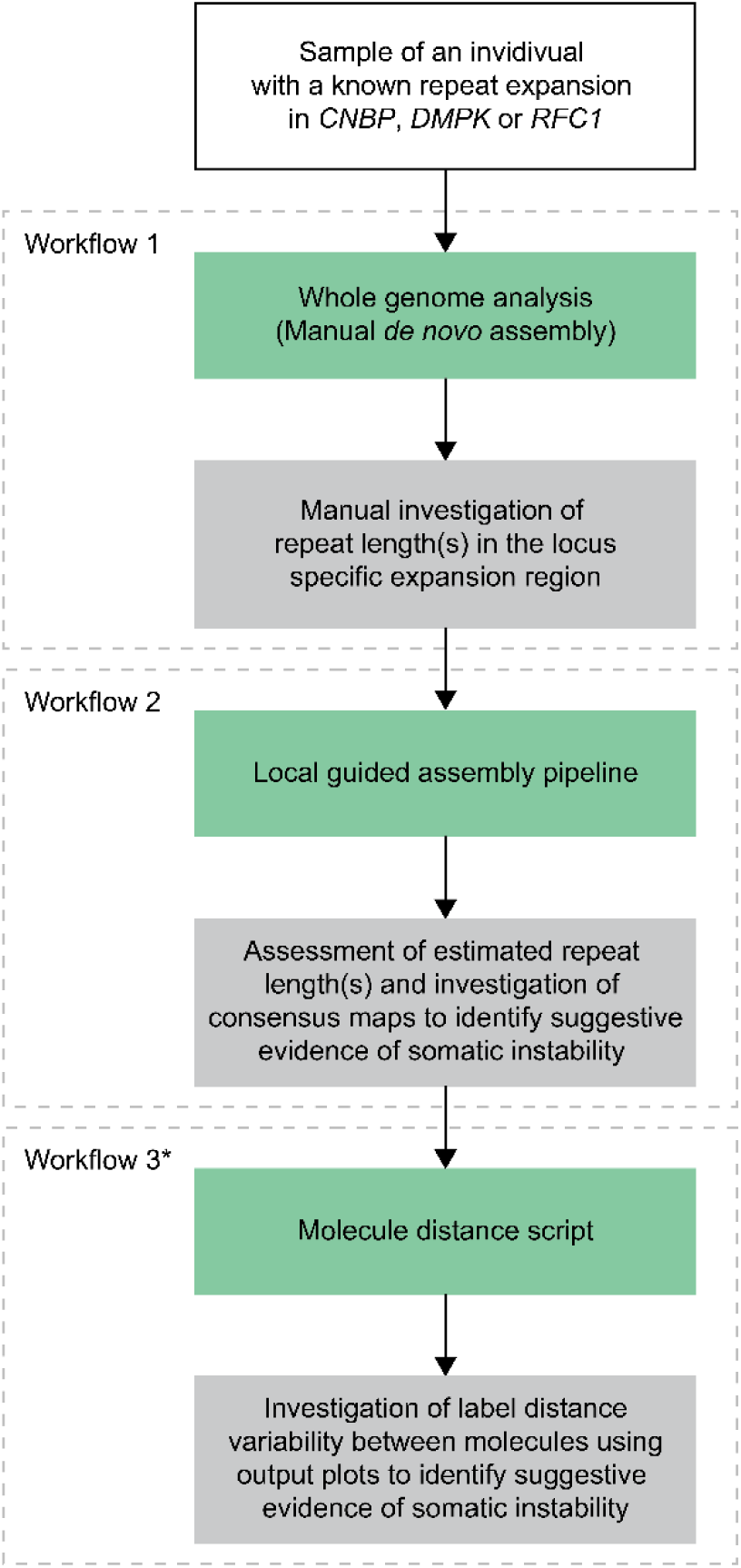
Total overview of the data analysis workflow. For each sample a *de novo* assembly was generated and the local guided assembly pipeline and molecule distance script were run. After each workflow the maps and/or molecules to calculate workflow specific repeat lengths were manually assessed. Green boxes denote the data analysis parts and gray boxes denote the data interpretation parts. (*) Workflow 1 and 2 were used to determine repeat lengths, while workflow 3 was used to identify potential somatic instability.

#### 1. Manual *de novo* assembly

In the manual *de novo* assembly workflow, for each individual a *de novo* assembly was generated on Solve 3.7.2 and Access 1.7.2 using default parameters against the GRCh38/hg38 reference genome. The *de novo* assembly was then used to estimate the repeat length for both alleles by calculating the genomic distance between the reference start and end label flanking the repeat locus of interest (**Fig. 2A** and **Supplemental Table S1**). The reference length between the two labels of interest was then subtracted from both allele lengths in the sample to get a repeat size estimate for both alleles. These sizes were then divided by the repeat unit length of the respective repeat locus to get the manual *de novo* assembly size estimates.

**Fig. 2:**
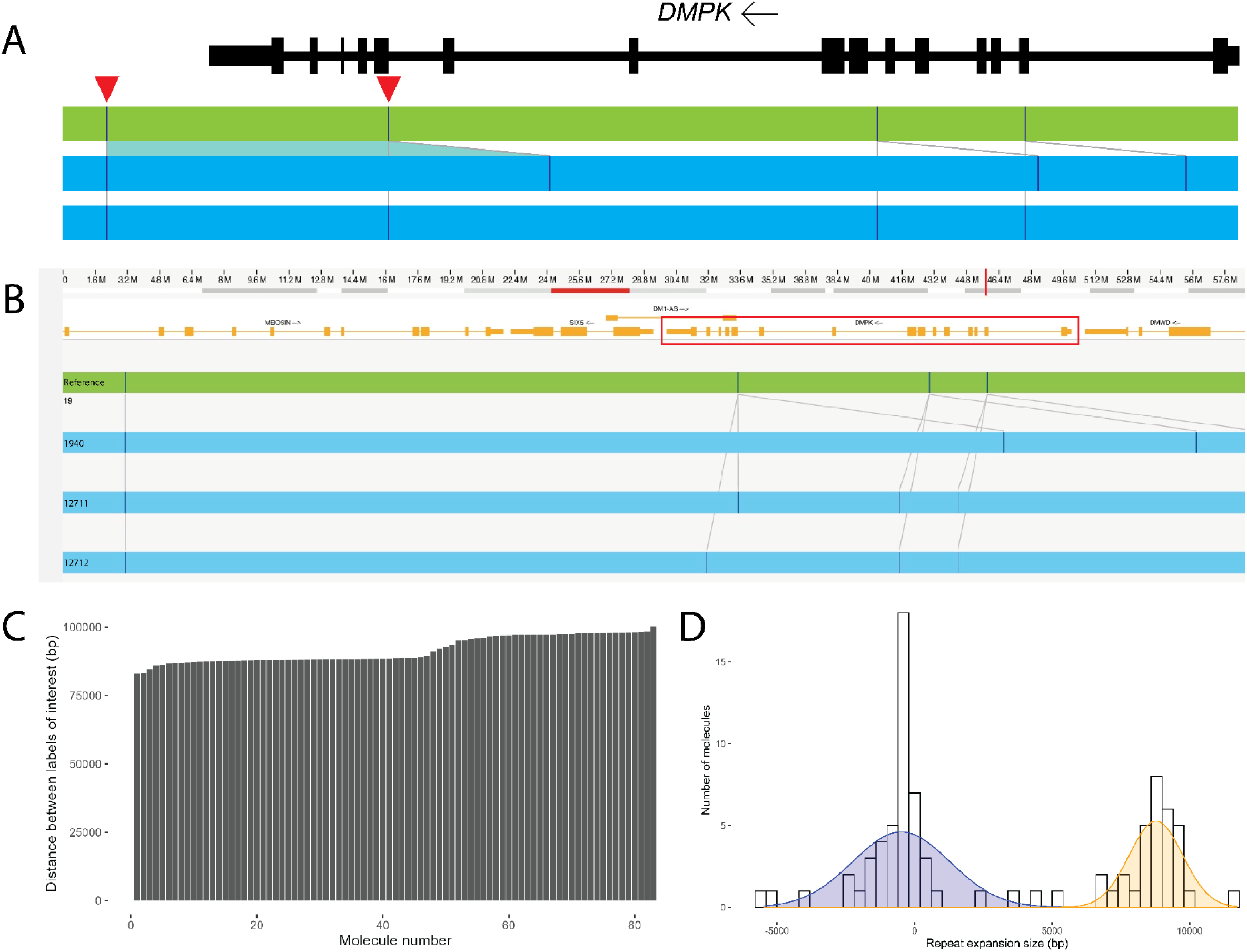
Overview of the data analysis outputs of the three OGM repeat expansion workflows for sample DMPK_10.This figure only shows the visual results of the data analysis. The results of the data interpretation are mainly the estimates of the actual repeat sizes resulting from the manual *de novo* assembly and local guided assembly workflows, as well as the visualization of the label distances in each molecule covering the locus of interest resulting from the molecule distance script. (A) Representation of the repeat expansion locus in the *de novo* assembly showing the position of the repeat expansion in the gene (3’UTR). Labels of interest are indicated by red arrow heads. These labels were used to manually calculate the repeat size by subtracting the reference distance (green bar) from the distances of the respective sample maps (blue bars). (B) Consensus guided assemblies across the *DMPK* repeat expansion locus. The *DMPK* gene is indicated by the red box. Based on the estimated repeat length, each map is assigned to allele 1 or allele 2 in order to separate the two alleles. Final repeat sizes are calculated by combining the repeat sizes of the maps assigned to the same allele (see also Methods). (C) This bar plot shows the distance between the labels of interest in each molecule ordered from smallest to largest. (D) This histogram shows the result of the molecule distance script that automatically assigns molecules to one of the alleles. The blue peak represents allele 1, while the orange peak represents allele 2. Both the bar plot and histogram can then be used to assess whether a sample contains evidence for somatic instability or not.

#### 2. Local guided assembly

For the local guided assembly (local-GA) workflow, the local-GA script was run on command line with locus specific seed and coordinate files using default settings (van der Sanden et al. 2024) (https://github.com/bionanogenomics/local_guided_assembly). Each of the output analysis reports lists the consensus map IDs (**Fig. 2B**) and calculated repeat expansion counts for each of those consensus maps. Maps were subsequently assigned to one of the two different alleles based on the estimated repeat counts. Generally, an output analysis report could contain maps with no or short repeat counts and maps with a large repeat counts. For homozygous and bi- allelic repeat expansions, the maps for both alleles could present large repeat counts. If the local- GA workflow resulted in a single consensus map and only one allele was expanded in the manual *de novo* assembly workflow for the same sample, the single local-GA consensus map was used as heterozygous call. If for both alleles were expanded in the manual *de novo* assembly workflow, the single map was used as homozygous call. For repeat report maps with ambiguous repeat counts, the global mean of repeat counts was used as a cutoff value to assign allele 1 or 2. Maps reported with “-1” repeat counts were excluded since the repeat counts could not be determined. Resulting repeat lengths were used as local guided assembly size estimates.

#### 3. Molecule distance script

The molecule distance script (https://github.com/bionanogenomics/molecule_distance) workflow was run on command line and required the intermediate *alignmolvref* files from the local- GA workflow. This *alignmolvref* result shows molecules aligned to the reference assembly (GRCh38/hg38). The script subsequently queried the distance between two predefined labels in each molecule (**Supplemental Table S2**). In order to successfully calculate the distance between the two labels of interest, only the molecules that contain both labels of interest were considered. Genomic distances were calculated using the distance between the start and end coordinates of the labels of interest in each molecule. Resulting repeat lengths were used as input for generating bar plots and histograms that visualize the repeat lengths to provide evidence for potential somatic instability (**Fig. 2C and 2D**)

### OGM repeat data interpretation

First, we determined for the manual *de novo* assembly workflow and local-GA workflow if a repeat expansion in the locus of interest of each respective sample was detected. A repeat was found to be detected when the result of the workflow identified that the longest allele was expanded beyond a gene specific repeat size threshold. For *CNBP* and *DMPK* the pathogenic repeat size threshold was used as gene specific threshold, while for *RFC1* a repeat size threshold of 20 repeat units was used (**Table 1**). Subsequently, for the *RFC1* samples, we assessed whether the results of the SOC corresponded with the results of the two OGM sizing workflows. For each detected *RFC1* repeat expansion we determined whether it was mono-allelic, bi-allelic or homozygous by comparing the detected repeat size(s) to the respective gene specific repeat size thresholds. The results of the two OGM workflows were then independently compared to the results of the SOC. Both OGM workflows had to indicate the same type of repeat as the SOC. If SOC reported a homozygous repeat expansion, OGM was allowed to identify both a homozygous and a bi-allelic repeat expansion. Finally, the actual repeat sizes resulting from the manual *de novo* assembly workflow and the local-GA workflow were compared to the repeat sizes reported after SOC. For each sample we determined whether at least one of the two OGM workflows identified a repeat expansion larger or equal to the SOC result.

### Detecting somatic instability

In order to identify potential somatic instability, multiple checks were performed. Firstly, the number of assembled maps at the region of interest in the local-GA data might indicate mosaicism (**Fig. 3A**). Stable repeat expansions usually form two maps during local-GA, indicating the reference and expanded allele. Additional maps are formed by molecules of unstable repeats clustered by the pipelines. Secondly, in the Bionano Access genome browser view, the molecule alignments to each of the assembled local-GA maps were visualized to search for a “gradient” of label distance in the molecule pile-up (**Fig. 3B**). Such a gradient might also indicate mosaicism. Finally, the molecule-to-reference alignment plots - or molecule distance plots - generated by the molecule distance script were examined for evidence of unstable alleles. When the expanded allele portion of a stable repeat locus is visualized using the molecule distance script, the molecule distances plateau at a certain length. Molecule distance bar plots with a steep gradient or a “stairway” distribution of label distances and histograms with a ‘smear’ instead of a peak, would suggest somatic instability (**Fig. 3C** and **3D**). We considered the data suggestive of somatic instability if a sample had both multiple consensus maps and a gradient distribution of molecule distances.

**Fig. 3:**
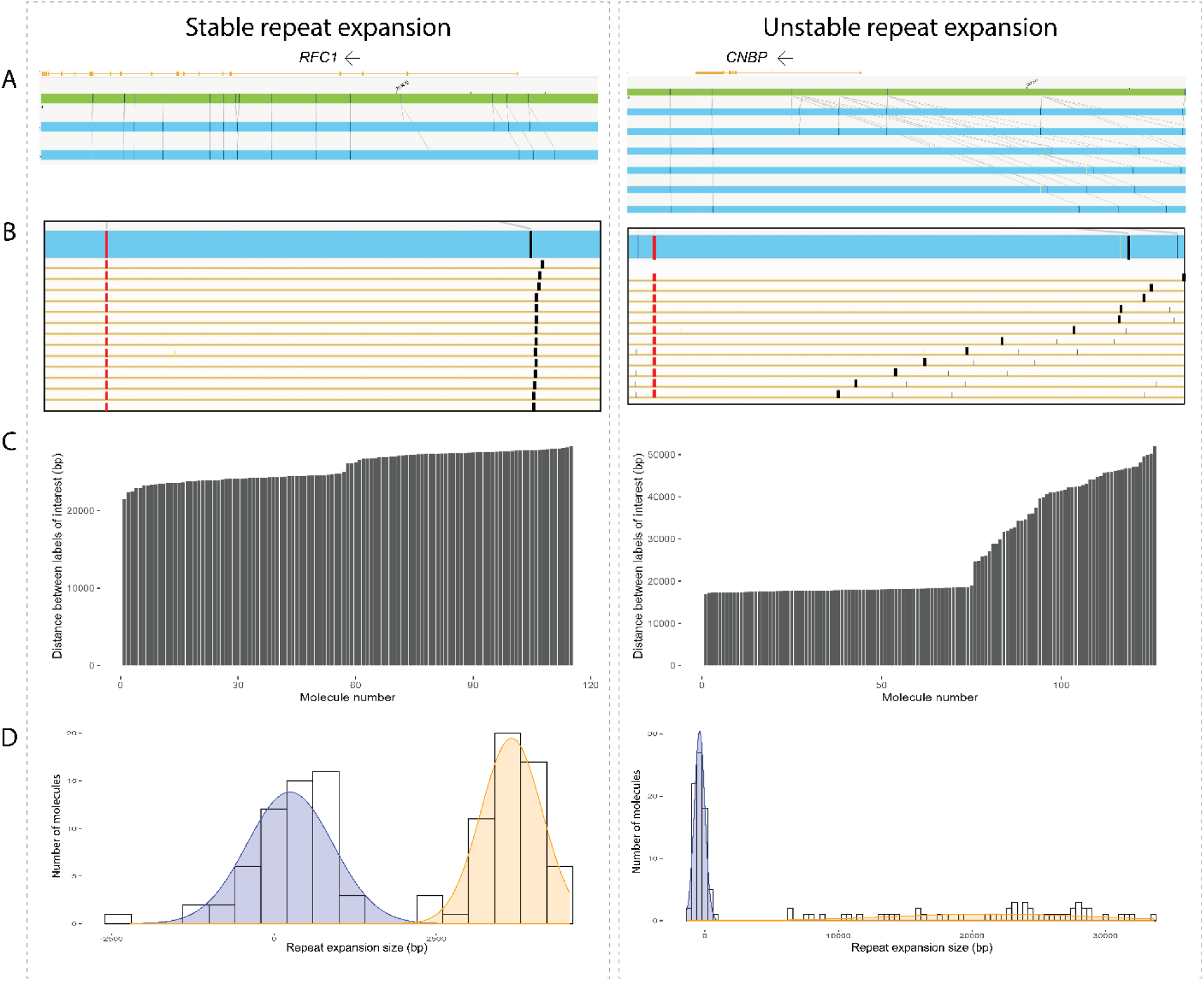
Representative plots of a sample with evidence and without evidence of somatic instability. The left part represents a stable *RFC1* repeat expansion and the right part represents a unstable *CNBP* repeat expansion. (A) The number of assembled maps at the region of interest in the local- GA data might indicate somatic instability. In this case the stable repeat had two consensus maps while the unstable repeat had six consensus maps. (B) A gradient of label distance in the molecule pile-up might also indicate mosaicism. The stable repeat had no gradient, while the unstable repeat presented a gradient of label distances based on the large variability in the distance between the red label and black label in each molecule. This variability results in the gradient or ‘stairway’ pattern. (C) The molecule distance script output plots show the repeat expansion size that is detected in each molecule by determining the distance between two specific labels of interest. This bar plot represents the distance between the labels of interest in each molecule ordered from smallest to largest. Molecule distance bar plots with a steep gradient or a stairway distribution of label distances would suggest somatic instability. The stable repeat had no stairway pattern, while the unstable repeat showed a stairway pattern for the expanded allele. The plot for the stable repeat visualizes the separation of the smaller allele and the larger allele around the middle of the plot (molecule number 57). The plot for the unstable repeat visualizes the same separation of the smaller allele and the larger allele (around molecule number 75). (D) The histogram plots outputted by the molecule distance script represent the separation of the two alleles based on the label distances in each molecule. The smaller alleles are indicated with blue peaks and the larger alleles are indicated with orange peaks. A ‘smear’ instead of a real peak in the histogram for one of the alleles might indicate somatic instability. For the stable repeat no smear was detected, while the unstable repeat presented with a ‘smear’ for the expanded allele. This is due to large variability in molecule label distances and therefore repeat expansion size.

## RESULTS

### Standard of care results

For all 85 individuals, SOC genetic testing previously identified at least a mono-allelic repeat expansion in *CNBP*, *DMPK* or *RFC1* that was larger than the pathogenic threshold (**Table 2**). All individuals with a mono-allelic repeat expansion in *DMPK* or *CNBP* resulted in the diagnosis of myotonic dystrophy type 1 or 2, respectively. Of the 30 samples with a repeat expansion in *DMPK,* 21 had a repeat expansion >150 units (450 bp) reported after SOC and based on this result we expected these repeat expansions to be larger than the formal structural variant detection limit of OGM, which is currently ∼500 bp. The nine remaining *DMPK* repeat expansions were determined to be smaller than 500 bp in size (range 61-159 units or 183-477 bp) and thereby below the formal OGM resolution cutoff. In the case of the individuals with a *RFC1* repeat expansion, 19 of the 30 individuals had a bi-allelic pathogenic AAGGG repeat expansion resulting in a diagnosis of cerebellar ataxia, neuropathy and vestibular areflexia syndrome (CANVAS), respectively. One other patient had a bi-allelic AAAAG repeat expansion that is considered to be benign. In addition, five other individuals were carriers of a pathogenic AAGGG repeat expansion of one allele, but carried a benign AAAAG or AAAGG repeat expansion on the other allele. The five remaining individuals were carriers of a mono-allelic AAGGG *RFC1* repeat expansion without an indication of a repeat expansion or other genetic variant on the other allele. The SOC had a detection threshold of >75 repeat units for *CNBP* and >150 repeat units for *DMPK*. For *RFC1* the SOC only predicted a mono- or bi-allelic repeat expansion, without providing any predictions of the expanded repeat size (**Table 2**).

**Table 2:** Sample and analysis overview. For each sample this table presents the SOC result, as well as the repeat size estimates from the two OGM sizing workflows (manual *de novo* assembly and local guided assembly) and the somatic instability assessment from the molecule distance script workflow. For *CNBP* and *DMPK* the table indicates whether the dominant repeat allele was detected (“Detected”). For *RFC1* we also checked whether OGM identified a mono-allelic, bi-allelic or homozygous repeat expansion. For the molecule distance script “A” denotes multiple consensus maps and “B” denotes a gradient in the molecule distances. We considered somatic instability in cases where both “A + B” provided suggestive evidence.

|  |  | SOC |  |  | Manual *de novo* assembly |  |  | Local guided assembly |  |  | Molecule distance script |
| --- | --- | --- | --- | --- | --- | --- | --- | --- | --- | --- | --- |
| Sample ID | Sample material | Allele 1 | Allele 2 | Conclusion | Allele 1 | Allele 2 | Conclusion | Allele 1 | Allele 2 | Conclusion | Somatic instability |
| CNBP_01 | EDTA blood | >75 | 18 | Detected | 3681 | 20 | Detected | 61 | 0 |  | - |
| CNBP_02 | EDTA blood | >75 | 15 | Detected | 6331 | 39 | Detected | 5000 | 10 | Detected | A + B |
| CNBP_03 | EDTA blood | >75 | 13 | Detected | 6517 | 53 | Detected | 3659 | 23 | Detected | A + B |
| CNBP_04 | EDTA blood | >75 | 15 | Detected | 7155 | -27 | Detected | 4401 | 4 | Detected | A + B |
| CNBP_05 | EDTA blood | >75 | 15 | Detected | 8042 | 0 | Detected | 3521 | 33 | Detected | A + B |
| CNBP_06 | EDTA blood | >75 | 15 | Detected | -14 | -15 |  | 3687 | 21 | Detected | A + B |
| CNBP_07 | EDTA blood | >75 | 16 | Detected | 5212 | -12 | Detected | 5000 | 10 | Detected | A + B |
| CNBP_08 | EDTA blood | >75 | 17 | Detected | 2502 | 2 | Detected | 2661 | 12 | Detected | A + B |
| CNBP_10 | EDTA blood | >75 | 13 | Detected | 2874 | -20 | Detected | 2963 | 11 | Detected | A + B |
| CNBP_11 | EDTA blood | >75 | 16 | Detected | 375 | 11 | Detected | 254 | 34 | Detected | A + B |
| CNBP_12 | EDTA blood | >75 | Normal | Detected | 3471 | -16 | Detected | 3346 | 14 | Detected | A |
| CNBP_13 | EDTA blood | >75 | 16 | Detected | 4634 | -14 | Detected | 4330 | 3 | Detected | A + B |
| CNBP_14 | EDTA blood | >75 | 16 | Detected | 5244 | -2 | Detected | 4186 | 49 | Detected | A + B |
| CNBP_15 | EDTA blood | >75 | Normal | Detected | 2183 | -1 | Detected | 2092 | 18 | Detected | A + B |
| CNBP_16 | EDTA blood | >75 | Normal | Detected | 3221 | 320 | Detected | 3201 | 0 | Detected | A + B |
| CNBP_17 | EDTA blood | >75 | 9 | Detected | 6275 | -13 | Detected | 5000 | 2 | Detected | A + B |
| CNBP_18 | EDTA blood | >75 | 15 | Detected | 1915 | -29 | Detected | 1656 | 15 | Detected | A + B |
| CNBP_19 | EDTA blood | >75 | Normal | Detected | 1460 | -2 | Detected | 1574 | 0 | Detected | A + B |
| CNBP_20 | EDTA blood | >75 | Normal | Detected | 3977 | -7 | Detected | 3577 | 11 | Detected | A + B |
| CNBP_21 | EDTA blood | >75 | 18 | Detected | 288 | 30 | Detected | 244 | 8 | Detected | A + B |
| CNBP_22 | EDTA blood | >75 | 17 | Detected | 1683 | -24 | Detected | 1725 | 70 | Detected | A + B |
| CNBP_23 | EDTA blood | >75 | 12 | Detected | 2131 | -25 | Detected | 2515 | 8 | Detected | A + B |
| CNBP_24 | EDTA blood | 134 | Normal | Detected | -13 | -14 |  | 3618 | 10 | Detected | A + B |
| CNBP_25 | EDTA blood | >75 | Normal | Detected | 1476 | -19 | Detected | 2626 | 9 | Detected | A + B |
| CNBP_26 | EDTA blood | >135 | Normal | Detected | 3737 | 14 | Detected | 3241 | 45 | Detected | A + B |
| DMPK_01 | EDTA blood | >150 | 11 | Detected | 269 | 55 | Detected | 247 | 35 | Detected | - |
| DMPK_02 | EDTA blood | >150 | 11 | Detected | 456 | 47 | Detected | 473 | 30 | Detected | - |
| DMPK_03 | EDTA blood | >150 | 5 | Detected | 252 | 57 | Detected | 116 |  | Detected | - |
| DMPK_04 | EDTA blood | 61 | 11 | Detected | 66 | 66 | Detected | 60 | 27 | Detected | B |
| DMPK_05 | EDTA blood | >150 | 5 | Detected | 457 | 54 | Detected | 485 | 30 | Detected | A + B |
| DMPK_06 | EDTA blood | 127 | 5 | Detected | 64 | 64 | Detected | 82 | 81 | Detected | B |
| DMPK_07 | EDTA blood | >150 | 12 | Detected | 378 | 28 | Detected | 358 | 20 | Detected | - |
| DMPK_08 | EDTA blood | 91 | 5 | Detected | 67 | 67 | Detected | 49 |  |  | B |
| DMPK_09 | EDTA blood | 96-130 | 5 | Detected | 71 | 71 | Detected | 68 |  | Detected | - |
| DMPK_10 | Cell pellet | >150 | 12 | Detected | 2829 | 37 | Detected | 2825 | 12 | Detected | A + B |
| DMPK_11 | Cell pellet | >150 | 5 | Detected | 233 | 58 | Detected | 231 | 12 | Detected | A + B |
| DMPK_12 | Cell pellet | >150 | 13 | Detected | 213 | 24 | Detected | 219 | 34 | Detected | B |
| DMPK_13 | Cell pellet | >150 | 13 | Detected | 163 | 21 | Detected | 167 | 33 | Detected | A + B |
| DMPK_14 | Cell pellet | >150 | 13 | Detected | 202 | 10 | Detected | 188 | 7 | Detected | B |
| DMPK_15 | Cell pellet | >150 | 6 | Detected | 1839 | 28 | Detected | 1768 | 53 | Detected | A + B |
| DMPK_16 | EDTA blood | >150 | 12 | Detected | 85 | 85 | Detected | 52 |  | Detected | B |
| DMPK_17 | EDTA blood | >150 | 5 | Detected | 491 | 41 | Detected | 510 | 15 | Detected | A + B |
| DMPK_18 | EDTA blood | >150 | 5 | Detected | 71 | 71 | Detected | 69 |  | Detected | B |
| DMPK_19 | EDTA blood | 73 | 12 | Detected | 54 | 54 | Detected | 61 | 0 | Detected | B |
| DMPK_20 | EDTA blood | >150 | 5 | Detected | 55 | 55 | Detected | 41 |  |  | B |
| DMPK_21 | EDTA blood | >150 | 12 | Detected | 1366 | 43 | Detected | 1347 | 23 | Detected | B |
| DMPK_22 | EDTA blood | 74 | 13 | Detected | 31 | 31 |  | 61 | 6 | Detected | A + B |
| DMPK_23 | EDTA blood | >150 | 7 | Detected | 372 | 17 | Detected | 369 | 25 | Detected | A + B |
| DMPK_24 | EDTA blood | 130 | 5 | Detected | 82 | 82 | Detected | 93 | 78 | Detected | A + B |
| DMPK_25 | EDTA blood | 159 | 5 | Detected | 79 | 79 | Detected | 109 | 63 | Detected | B |
| DMPK_26 | EDTA blood | 88 | 13 | Detected | 45 | 45 |  | 29 | 0 |  | A |
| DMPK_27 | EDTA blood | >150 | 5 | Detected | 1648 | 21 | Detected | 20 | 12 |  | B |
| DMPK_28 | EDTA blood | >150 | 14 | Detected | 440 | 41 | Detected | 393 | 3 | Detected | B |
| DMPK_29 | EDTA blood | >150 | 12 | Detected | 320 | 20 | Detected | 310 | 12 | Detected | B |
| DMPK_30 | EDTA blood | >150 | 29 | Detected | 290 | 63 | Detected | 131 |  | Detected | B |
| RFC1_01 | EDTA blood | >>AAGGG | >>AAGGG | Homozygous | 1487 | 1174 | Bi-allelic | 1497 | 1160 | Bi-allelic | A |
| RFC1_02 | EDTA blood | >>AAGGG | >>AAAAG | Bi-allelic | 738 | 452 | Bi-allelic | 751 | 458 | Bi-allelic | - |
| RFC1_03 | EDTA blood | >>AAGGG | >>AAAAG | Bi-allelic | 883 | 111 | Bi-allelic | 897 | 126 | Bi-allelic | A |
| RFC1_04 | EDTA blood | >>AAGGG | >>AAAAG | Bi-allelic | 750 | 97 | Bi-allelic | 760 | 99 | Bi-allelic | - |
| RFC1_05 | EDTA blood | >>AAGGG | 11 AAAAG | Mono-allelic | 1167 | -5 | Mono-allelic | 1175 | 4 | Mono-allelic | A + B |
| RFC1_06 | EDTA blood | >>AAAAG | >>AAAAG | Homozygous | 1565 | 1278 | Bi-allelic | 1579 | 1283 | Bi-allelic | A |
| RFC1_07 | EDTA blood | >>AAGGG | 11 AAAAG | Mono-allelic | 625 | -3 | Mono-allelic | 643 | 10 | Mono-allelic | - |
| RFC1_08 | EDTA blood | >>AAGGG | >>AAGGG | Homozygous | 840 | 840 | Homozygous | 856 | 856 | Homozygous | - |
| RFC1_09 | EDTA blood | >>AAGGG | >>AAGGG | Homozygous | 1506 | 770 | Bi-allelic | 1499 | 778 | Bi-allelic | A |
| RFC1_10 | EDTA blood | >>AAGGG | >>AAGGG | Homozygous | 1289 | 1175 | Bi-allelic | 1307 | 1131 | Bi-allelic | A |
| RFC1_11 | EDTA blood | >>AAGGG | >>AAGGG | Homozygous | 812 | 812 | Homozygous | 818 | 818 | Homozygous | - |
| RFC1_12 | EDTA blood | >>AAGGG | >>AAGGG | Homozygous | 1106 | 895 | Bi-allelic | 1121 | 888 | Bi-allelic | A |
| RFC1_13 | EDTA blood | >>AAGGG | >>AAGGG | Homozygous | 927 | 927 | Homozygous | 933 | 933 | Homozygous | - |
| RFC1_14 | EDTA blood | >>AAGGG | >>AAGGG | Homozygous | 725 | 725 | Homozygous | 740 | 737 | Bi-allelic | - |
| RFC1_15 | EDTA blood | >>AAGGG | >>AAGGG | Homozygous | 873 | 811 | Bi-allelic | 887 | 783 | Bi-allelic | A |
| RFC1_16 | EDTA blood | >>AAGGG | 11 AAAAG | Mono-allelic | 1104 | -11 | Mono-allelic | 1097 | 4 | Mono-allelic | A + B |
| RFC1_17 | EDTA blood | >>AAGGG | 9 AAAAG | Mono-allelic | 711 | -1 | Mono-allelic | 723 | 8 | Mono-allelic | B |
| RFC1_18 | EDTA blood | >>AAGGG | >>AAGGG | Homozygous | 1444 | 1134 | Bi-allelic | 1474 | 1127 | Bi-allelic | A + B |
| RFC1_19 | EDTA blood | >>AAGGG | >>AAGGG | Homozygous | 223 | 40 | Bi-allelic | 235 | 51 | Bi-allelic | A |
| RFC1_20 | EDTA blood | >>AAGGG | >>AAAAG | Bi-allelic | 494 | 100 | Bi-allelic | 502 | 106 | Bi-allelic | - |
| RFC1_21 | EDTA blood | >>AAGGG | >>AAGGG | Homozygous | 703 | 600 | Bi-allelic | 715 | 600 | Bi-allelic | A + B |
| RFC1_22 | EDTA blood | >>AAGGG | >>AAGGG | Homozygous | 1161 | 905 | Bi-allelic | 1180 | 911 | Bi-allelic | A |
| RFC1_23 | EDTA blood | >>AAGGG | >>AAGGG | Homozygous | 1028 | 701 | Bi-allelic | 1054 | 733 | Bi-allelic | - |
| RFC1_24 | EDTA blood | >>AAGGG | 11 AAAAG | Mono-allelic | 602 | 2 | Mono-allelic | 615 | 11 | Mono-allelic | A |
| RFC1_25 | EDTA blood | >>AAGGG | >>AAGGG | Homozygous | 855 | 854 | Bi-allelic | 868 | 868 | Homozygous | - |
| RFC1_26 | EDTA blood | >>AAGGG | >>AAGGG | Homozygous | 714 | 573 | Bi-allelic | 734 | 585 | Bi-allelic | - |
| RFC1_27 | EDTA blood | >>AAGGG | >>AAGGG | Homozygous | 973 | 973 | Homozygous | 987 | 403 | Bi-allelic | A |
| RFC1_28 | EDTA blood | >>AAGGG | >>AAGGG | Homozygous | 875 | 875 | Homozygous | 893 | 893 | Homozygous | - |
| RFC1_29 | EDTA blood | >>AAGGG | >>AAGGG | Homozygous | 1071 | 890 | Bi-allelic | 1073 | 905 | Bi-allelic | - |
| RFC1_30 | EDTA blood | >>AAGGG | >>AAAAG | Bi-allelic | 743 | 74 | Bi-allelic | 754 | 85 | Bi-allelic | A |

### Detecting repeat expansions using optical genome mapping

The OGM approach consisted of the generally available *de novo* assembly pipeline as well as two workflows that were developed as part of this study, *i.e.* local guided assembly (local-GA) and molecule distance script. In this study, we used these three different and complementary analytical workflows based on the OGM BNX molecule files to either estimate the size of both alleles at the respective locus of interest (manual *de novo* assembly and local-GA) or to assess the somatic stability of the detected repeat expansion(s) (molecule distance script) (**Fig. 1**). The manual *de novo* assembly workflow identified a repeat expansion beyond the gene specific repeat size threshold in 81/85 (95.3%) samples, while the local-GA identified a repeat expansion beyond the gene specific repeat size threshold in 80/85 (94.1%) samples (**Table 2** and **Supplemental Fig. S1**). Jointly, we were able to identify a repeat expansion for 84 of the 85 samples by combining the results of the two different sizing workflows, even when considering the expected expansions smaller than the 500 bp formal cutoff for SV calling with OGM. The one remaining sample (DMPK_26) had a repeat size of 88 repeat units based on SOC, but only a premutation was suggested by the OGM findings with 45 repeat units called by the manual *de novo* assembly. Of the 84 detected repeat expansions, 77 were called by both workflows and the remaining seven were called as repeat expansion by one of the two workflows (**Supplemental Fig. S1** and **Table 2**). Remarkably this even included eight samples with *DMPK* repeat expansion lengths <500 bp, the formal detection limit of OGM. Of the latter, six were called by both sizing OGM workflows, while the other two were only called by one of the two workflows.

### Concordance between OGM and SOC

Myotonic dystrophy type 1 and 2 are both autosomal dominant disorders, which is why we only expected heterozygous repeat expansions in the *DMPK* and *CNBP* samples. For all these samples except one, OGM identified the heterozygous repeat expansion. However, CANVAS is an autosomal recessive disorder caused by compound heterozygous or homozygous repeat expansions in *RFC1*, which required to assess the repeat length in both alleles. Therefore, we confirmed whether both OGM workflows resulted in the same type of repeat expansion as reported after SOC, *i.e.* a mono-allelic, bi-allelic or homozygous repeat expansion and for all 30 *RFC1* samples, OGM confirmed the SOC results (**Table 2**).

In addition, the actual repeat lengths of the two OGM workflows (manual *de novo* assembly and local-GA) were compared to the repeat lengths reported after SOC. For all 25 *CNBP* and 30 *RFC1* samples the repeat lengths identified by OGM had at least the length reported after SOC (**Table 2**) and these results were considered concordant. In the case of *DMPK*, for 20 of the 30 samples the repeat expansion lengths were also concordant with SOC, while for the other ten samples OGM presented different calls for the absolute repeat length compared to the SOC (**Table 3**). For seven of these ten samples, the SOC identified a repeat expansion length <500 bp, the formal resolution limit of OGM. The remaining three samples had an expected repeat length >500 bp (based on SOC). The results for *DMPK* also indicated that the manual *de novo* assembly overestimated the repeat size of the expected wildtype allele (based on SOC) to be ≥50 repeat units (range 54-85 repeat units or 162-255 bp) for 15 samples. All but three of these wildtype alleles were called <50 repeat units by the local-GA workflow, suggesting that the local-GA may be more accurate in distinguishing wildtype and small repeat expansions.

**Table 3:** Overview of repeat expansions with different calls for absolute repeat size.

| Sample ID | SOC |  | Manual *de novo* assembly |  | Local guided assembly |  |
| --- | --- | --- | --- | --- | --- | --- |
|  | Allele 1 | Allele 2 | Allele 1 | Allele 2 | Allele 1 | Allele 2 |
| DMPK_06 | 127 | 5 | 64 | 64 | 82 | 81 |
| DMPK_08 | 91 | 5 | 67 | 67 | 49 | wt |
| DMPK_09 | 96-130 | 5 | 71 | 71 | 68 | wt |
| DMPK_16 | >150 | 12 | 85 | 85 | 52 | wt |
| DMPK_18 | >150 | 5 | 71 | 71 | 69 | wt |
| DMPK_19 | 73 | 12 | 54 | 54 | 61 | 0 |
| DMPK_20 | >150 | 5 | 55 | 55 | 41 | wt |
| DMPK_22 | 74 | 13 | 31 | 31 | 61 | 6 |
| DMPK_24 | 130 | 5 | 82 | 82 | 93 | 78 |
| DMPK_25 | 159 | 5 | 79 | 79 | 109 | 63 |

### Distinguishing between the two repeat alleles in bi-allelic repeats

OGM also allowed to distinguish between the two *RFC1* repeat expansion alleles of similar size for 19/25 *RFC1* repeat expansion samples for which SOC identified a bi-allelic or homozygous expansion. For the remaining 6 RFC1 repeat expansion samples, OGM detected a homozygous repeat expansion, which confirmed the SOC results (**Table 2**).

### Comparing the exact repeat sizes across the two OGM sizing pathways

One of the advantages of OGM over the SOC is that it also provided estimates of the actual size of large repeats starting from ∼500 bp in size. This allowed us to compare the repeat size estimates of each sample across the two OGM repeat sizing workflows. The ranges of the detected repeat expansions detected by both sizing workflows were [288 – 8042] and [244 – 5000] for *CNBP*, [54 – 2829] and [52 – 2825] for *DMPK*, and [223 – 1565] and [235 – 1579] for *RFC1* for the manual *de novo* assembly workflow and local-GA workflow respectively (**Table 2**). There was a strong, significant correlation among the manual *de novo* assembly and the guided assembly workflows (R=0.94, P = <0.001) (**Fig. 4**). The intercept for the comparison was 292, indicating a deviation between the two workflows, which was confirmed by the fact that the manual *de novo* assembly workflow overestimated the size of the repeat expansion alleles by an average of 3%.

**Fig. 4:**
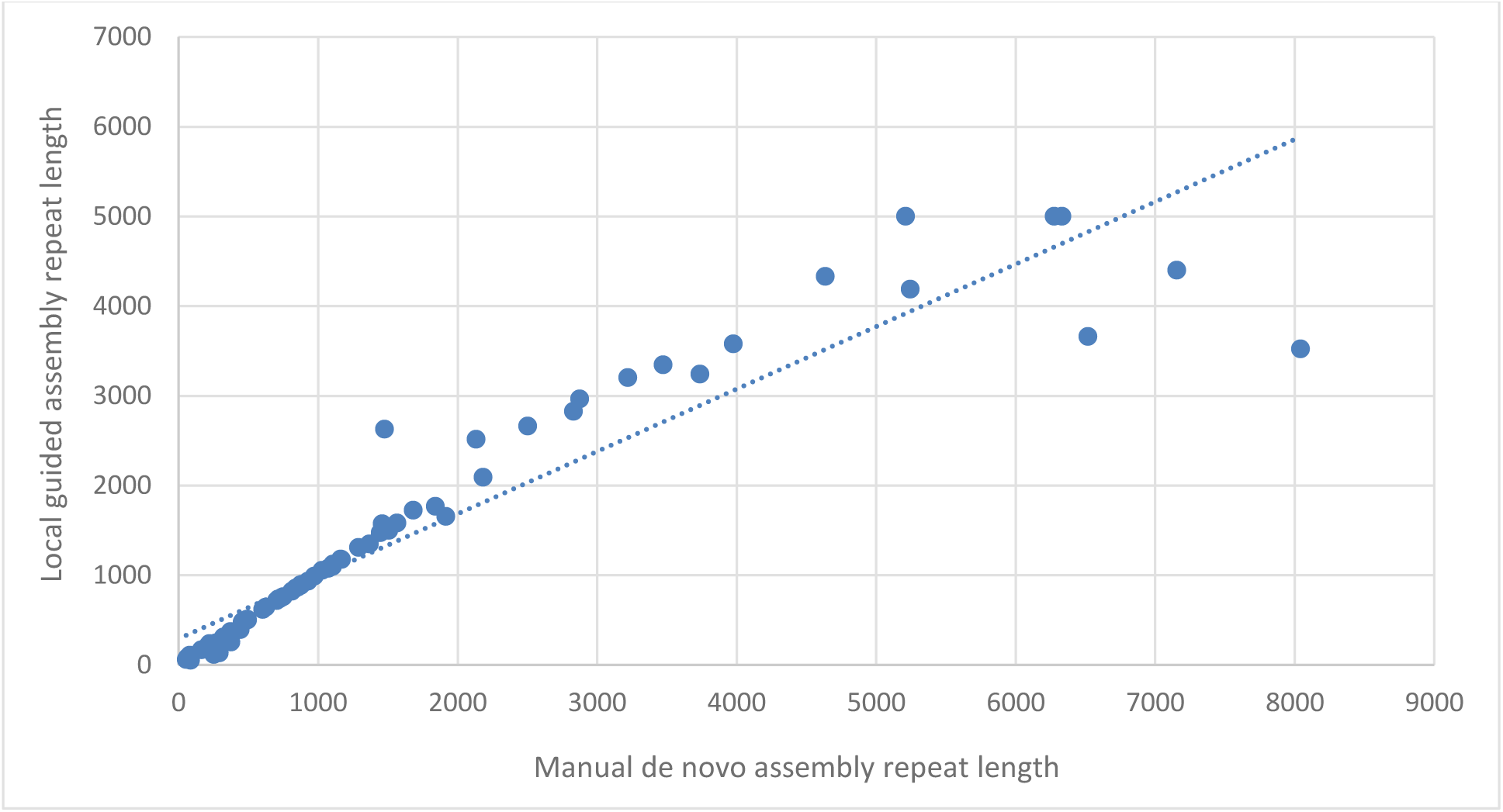
Correlation between the manual *de novo* assembly repeat lengths and the local guided assembly repeat lengths. For this correlation assessment we only used the 77/85 (90.6%) samples for which both the manual *de novo* assembly workflow and the local-GA workflow detected a repeat expansion.

### Detecting somatic instability

Based on the number of consensus maps and corresponding molecules resulting from the local- GA workflow and the visual inspection of the bar plot and histogram resulting from the molecule distance script workflow (**Fig. 3**), we detected suggestive evidence of somatic instability in 36/85 samples. Of these, 23 were *CNBP* samples, 9 *DMPK* samples and 4 *RFC1* samples. Remarkably, of the 25 samples with the largest repeat alleles (>1,500 repeat units), only two had no suggestive evidence of instability. It seems that the molecular distance script workflow may be best suited to detect instability, and 16 different samples show a suggestive pattern for instability by this tool alone. Due to the suspected somatic instability, the estimated repeat sizes may vary more than the estimated repeat sizes of samples without a somatic instability suspicion. This result suggests the benefit of sequentially using the different OGM repeat expansion workflows, especially in the case of samples that are suspected to present with somatic instability.

## DISCUSSION

Determining the exact length of specific repeat expansions is of great importance for the patient and its family due to the rough correlation between repeat size and disease severity and age of onset, but also due to genetic anticipation. Unfortunately, current molecular diagnostic efforts for repeat expansion disorders entail labor-intensive and time-consuming PCR and / or Southern blot efforts. The current SOC only determines a repeat size range but does not detect/estimate the actual repeat length (due to artifacts or resolution limits of the respective tests). Also the read size of short-read sequencing methods have proven to be too limited to accurately detect all repeat expansions and long-read sequencing is still not routinely used in most laboratories and is currently too expensive, while it allows the detection of an increasing amount of novel repeat expansion and contraction disorders most recently (Pellerin et al. 2023). Here, we present a generic assay that works for three different repeat loci (*i.e. CNBP*, *DMPK* and *RFC1*) and most likely also for all other repeat expansion loci for which the pathogenic repeat size extends beyond ∼300 bp in size. We utilize that OGM uses native DNA molecules without any experimental noise (*e.g.* PCR artifacts, or bias for one of the alleles), which allows to detect very long repeat expansions. An additional benefit of this approach is the possibility to detect somatic instability in the repeat expansion of interest.

Overall, our results increased the repeat allele sizing resolution for all 84 of the 85 investigated repeat expansion samples. Being able to provide a more accurate repeat length measurement, especially for very long repeat expansion alleles, is one of the apparent strengths of OGM. Here, we even detected *CNBP* expansions >5,000 repeat units, suggesting OGM has no upper size limit, which may still exist for most short- and long-read sequencing approaches. In addition, for 19 of the *RFC1* samples, the SOC reported a bi-allelic or homozygous repeat expansion and OGM allowed to distinguish between the two alleles of similar size, which is not possible with current SOC. With OGM enabling to confirm, size and distinguish both heterozygous and bi-allelic repeat expansions, it also increases molecular diagnostic capabilities and allows for improved patient and family counseling. This is particularly important for families with *RFC1* repeat expansions because the repeat length of these expansion alleles, and especially the length of the smaller allele, is an important factor for predicting disease onset, phenotype variability and severity (Currò et al. 2024).

An additional benefit of this approach is the possibility to detect somatic instability in the repeat expansion of interest, a phenomenon that could potentially lead to variability in disease severity and age of onset, especially if affected tissue could be sampled (Monckton et al. 1995; Wong et al. 1995; Gomes-Pereira et al. 2004; Swami et al. 2009; Goold et al. 2021). Here, we detected evidence of somatic instability for at least 36/85 samples or 30.0% of *DMPK* samples, 92.0% of *CNBP* samples and 16.0% of *RFC1* samples. Strikingly, instability seems to occur for almost all long repeats, *i.e.* all but two of the 25 largest repeats. Finding this large number of somatically unstable repeat expansions was not expected beforehand. For *CNBP* (Alfano et al. 2022) and *DMPK* (Morales et al. 2022) repeat expansion alleles the presence of somatic instability is well known, however so far *RFC1* repeat alleles have been considered stable and evidence for somatic instability in *RFC1* repeat expansions is limited (Currò et al. 2024). Finding this new evidence, highlights an opportunity for future repeat expansion research using OGM, as OGM can easily identify somatic instability for various repeat loci. Sensitivity of this approach may go up with generating higher coverage with OGM. Also updates of the molecular distance script workflow shall allow to determine more accurate cut-offs for instability in the future.

Notwithstanding the accurate repeat expansion detection and improved allele sizing resolution using OGM, our results confirm the suspicion that OGM might not be accurate for repeat sizes smaller than 500 bp. For ten *DMPK* cases smaller repeat sizes than expected by SOC were detected. For seven of those, also SOC confirmed allele sizes smaller than 500 bp. However, due to technical difficulties such as extinction of the repeat-primed PCR signal, the precision of SOC may also not represent the ground truth in all cases. Our data also suggests that using a pathogenic repeat length threshold >300 bp (74 repeat units for CNBP *i.e.* 296 bp) does not result in false positive findings *i.e.* the overestimation of wildtype alleles, while a smaller threshold (50 repeat units for DMPK *i.e.* 150 bp) may result in an overestimation of wildtype allele sizes as seen for 15/85 samples (all called with >50 repeat units for the suspected wildtype *DMPK* allele). This overestimation of wildtype allele sizes seems to mainly occur in the manual *de novo* assembly workflow and is less of an issue when using the local-GA workflow. In total, our study suggests that OGM is highly accurate for identifying large repeat expansions. There was only one sample for which full expansion status was called by SOC but only premutation status by OGM. This may not surprise as this sample presented with a repeat size by SOC of only 88 repeat units or 264 bp.

Even though both the manual *de novo* assembly workflow and the local-GA workflow use the same BNX molecule file as starting input, we show that there is a deviation between the repeat sizes as estimated by these two different workflows (**Fig. 4** and **Table 2**). The correlation between the two workflows was highly significant, indicating that such a deviation is probably caused by a systematic error in one of the OGM repeat detection workflows, which was quantified as an average 3% overestimation of the repeat sizes by the manual *de novo* assembly workflow compared to the local-GA workflow. However, it seems that this overestimation is mainly caused by seven large *CNBP* repeats that are overestimated by more than 20% in the manual *de novo* assembly compared to the local guided assembly (**Fig. 4** and **Table 2**). Further validation processes (*e.g.* using targeted long-read sequencing), could help to clarify which of the two workflows provides the most accurate size estimation. However, our results also suggest that it may not be necessary to choose only one of the three OGM repeat workflows, because they can also be used sequentially or in parallel, which would create one single method for repeat expansion detection using OGM data. By using this single method, the different analysis workflows work together and can even complement each other. First, the manual *de novo* workflow can indicate a potential repeat expansion even beyond the currently specified 500 bp resolution cutoff of SV calling using OGM. Next, the local-GA allows a more targeted size estimate by collecting molecules and aligning these molecules to each other to create a consensus map for only the specific region of interest. The algorithm then determines the size of the expansion in the different maps specifically at the respective locus of interest. Finally, the molecule distance script can separate the two alleles and clearly visualize this separation by plotting individual molecule lengths at the locus of interest. The plots resulting from this latter part are particularly useful for identifying unstable repeat expansions. Altogether, this suggests that the three separate workflows work best in a complementary fashion and all three can be performed locally. The manual *de novo* assembly workflow can be performed using the Bionano Access analysis software by loading in a pre-generated *de novo* assembly file and the local-GA and molecule distance script workflows were developed as part of this study and are publicly available (https://github.com/bionanogenomics/local_guided_assembly/ and https://github.com/bionanogenomics/molecule_distance/) (van der Sanden et al. 2024).

The local-GA data not only provides repeat size estimates for both alleles, but it also generates confidence intervals for each repeat length. In this study these confidence intervals remained outside of the scope because we only worked with the repeat size estimates that could be equally compared between the two sizing workflows. However, being able to use these confidence intervals could be a very nice add-on for clinical labs when using OGM for repeat expansion detection, because the two different workflows present different repeat sizes and it may be difficult to rationalize which sizes to use. In addition, these confidence intervals may help to control for a potential systematic error in one of the OGM repeat detection workflows. However, potential somatic instability must be taken into account when using the confidence intervals, which suggests that potential somatic repeat instability should be assessed using the molecule distance script before using the confidence intervals in downstream analyses. In addition, the molecule distance script can be improved in identifying and characterizing somatic expansion alleles by implementing a statistical method to automate the output interpretation. Now the somatic instability assessment relies on manual inspection, but an automated model would help to reduce variability in reporting results.

Advantages of OGM over SOC and sequencing methods are not limited to the sizing resolution and detection of somatic instability. Considering the unexpectedly high level of somatic instability, OGM presents with another advantage, that is that higher coverage than for genome sequencing can routinely be reached with latest OGM iterations allowing coverage up to 1,500-fold without extra cost (Smith et al. 2023). In addition, OGM only uses natural UHMW DNA molecules that are not sheared and are not subjected to any obvious bias, such as PCR or sequencing bias. Even though the lab process for OGM requires up to 5 hours of hands-on time and contains multiple incubation steps, this method provides higher accuracy and higher throughput. Moreover, after analyzing the labeled DNA on the Saphyr® machine, the results can easily be reanalyzed for different repeat expansion loci, without the need to rerun any sample, while for SOC new PCRs or blots have to be performed. Since some repeat expansion disorders have overlapping phenotypic characteristics and strong heterogeneity of symptoms, this option of analyzing the entire human genome at once, proves a large benefit – and would allow OGM to become a truly generic test for all established expansion disorders for which expansions lead to SVs >500bp or even ∼300 bp as shown for the smallest alleles here (DMPK_04). In line with this, 11 additional samples with a repeat expansion in *ATXN10* (Morato Torres et al*. 2022)*, *C9orf72* (Barseghyan et al. 2022), *FXN*, *NOP56* or *STARD7* were also analyzed successfully (**Supplemental Table S3**), suggesting indeed that OGM is suited for most known repeat expansion disorders. Finally, if a repeat expansion disorder is suspected, but is not confirmed by OGM, the generated *de novo* assembly still allows to identify different types of SVs including other insertions and deletions, but also deletions, inversions and translocations. Hereby, this method is more versatile than other repeat expansion disorder tests in the SOC.

Besides the advantages of OGM over SOC and sequencing efforts, it also has a known limitation, being the inability to provide sequence context for all its SV calls and therefore also for the repeat expansion insertion calls. For certain repeat expansion disorders the sequence context can be of high importance, since repeat interruptions may cause repeat (in)stability and thereby mitigating the disease severity. Also for *RFC1* repeats where pathogenic AAGGG and normal AAAGG and AAGGG repeats are known, OGM cannot determine which type of repeat expansion is detected. Therefore, if the sequence context is of importance for the specific repeat expansion disorder, the OGM test still must be complemented with preferably (targeted) long-read sequencing, which adds to the financial considerations that have to be made before choosing OGM as the technology to detect those specific repeat expansions. In addition, a separate copy number variant or SV in or around the region of interest, as well as variation of the label-site can influence the results of the workflows. Therefore, a thorough inspection of the *de novo* assembly in workflow 1 using the circos plot or genome browser in the Bionano Access software is of great importance. When there is any indication of another large variant, the results of the different repeat detection workflows must be analyzed with extra care to prevent reporting of false positive or false negative results.

In conclusion, our data demonstrate that OGM can efficiently and accurately identify the repeat lengths across multiple STR loci simultaneously, thereby detecting large STR expansions and determining their absolute repeat sizes. This supports the technical validity of OGM for the detection of repeat expansion alleles larger than ∼300 bp in size. OGM increased the allele sizing resolution for 84/85 repeat samples, and it indicated 36 samples with suggestive evidence of somatic repeat instability. Our results also suggest that OGM can detect all large repeat expansions >300 bp in size using a single test, which is in contrast to current SOC that uses multiple gene specific tests to reach the same conclusions while potentially taking more time and being more expensive. This proves that OGM could serve as a more efficient workflow for repeat expansion detection. However, whether this increased efficiency can compensate for the unavailability of exact sequence context remains to be determined.

## DATA ACCESS

All data obtained of relevance to support the conclusions are presented in the manuscript or in the supplementary datafiles, for which more details are available upon reasonable request from the authors.

Local guided assembly script and accompanying files are available at: https://github.com/bionanogenomics/local_guided_assembly/blob/master/run_local_guided_assembly.sh

Molecule distance script is available at: https://github.com/bionanogenomics/molecule_distance/

## ACKNOWLEDGEMENTS

We would like to acknowledge colleagues from the diagnostic division of the Radboudumc (Genome Diagnostics Nijmegen) as well as the Radboud Genomics Technology Center for their support.

Dr. Hoischen was supported by the Solve-RD project. The Solve-RD project has received funding from the European Union’s Horizon 2020 research and innovation program under grant agreement No. 779257. This research was part of the Netherlands X-omics Initiative and partially funded by NWO (Dutch Research Council, 184.034.019).

## AUTHOR CONTRIBUTIONS

Conceptualization: EJK, AHo; Data curation: BvdS, KN, SS, MDG, JL, AWCP; Formal analysis: BvdS, KN, SS, MDG, JL, MP, RvB, MO, EKB, EK, AWCP; Funding acquisition: AHo; Investigation: KN, SS, MDG, JL, MP, RvB, MO, EKB, EK, AWCP; Methodology: BvdS, SS, MDG, JL, SLB, AWCP, AHa; Project administration: BvdS, EJK, AHo; Resources: AT, NV, IES, JG, MAC, AHa, AHo; Software: SS, MDG, JL, AWCP, AHa; Supervision: LELMV, AHa, EJK, AHo; Validation: BvdS, SS, MDG, JL, AWCP; Visualization: BvdS, SS, SLB, AWCP; Writing-original draft: BvdS, KN, EJK, AHo; Writing-review & editing: BvdS, KN, SS, MDG, JL, SLB, AT, NV, AWCP, AHo. All authors have contributed to the manuscript and have read and approved the final version of the manuscript.

